# Comparative assessment of mosquito size and receptiveness to *P. falciparum* infection of two rearing procedures for *Anopheles* mosquitoes

**DOI:** 10.1101/2023.03.09.531859

**Authors:** Wouter Graumans, Kjerstin Lanke, Marga van de Vegte-Bolmer, Rianne Stoter, Giulia Costa, Elena A Levashina, Annie S P Yang, Geert-Jan van Gemert, Teun Bousema

## Abstract

Malaria is transmitted when *Anopheline* mosquitoes ingest *Plasmodium* parasites during blood-feeding. Artificial feeding assays allow mosquitoes to take up blood from membrane feeders, and are widely used to study malaria transmission. These assays require large quantities of mosquitoes; insectaries optimize their rearing procedures to generate high yields of permissive, homogeneous mosquito populations. Rearing of *Anopheline stephensi* mosquitoes was protocolized at the Radboudumc in the 1980s, yet infection outcomes remain heterogeneous. This study explores possible improvements in mosquito rearing to improve homogeneity of the resulting mosquito populations. It compares the current mass-rearing standard with an adapted alternative approach from another institute that optimizes larval density per tray and applies a diet that has previously been reported. Differences between procedures were assessed by measuring mosquito size by proxy of wing-length and imbibed blood meal volume. To assess receptiveness to *P. falciparum* infection mosquitoes were fed cultured gametocytes. We observed a slight decrease in mosquito size when applying the alternative rearing procedure, but generated equally parasite-receptive *P. falciparum* mosquitoes compared to the standard procedure. We conclude that both rearing protocols can be used to generate susceptible mosquitoes for conducting malaria research.

## Background

Malaria, caused by *Plasmodium* parasites that are transmitted by infected *Anopheline* mosquitoes, remains the most important vector-borne disease worldwide [1]. Artificial mosquito feeding assays are important tools to test novel interventions against malaria, understand factors that determine vector competence and gain a better understanding of the infectiousness of natural malaria-infected carriers to mosquito vector [2]. Conducting these assays requires the rearing of infection-permissive mosquitoes that also successfully feed on membranes of artificial feeding apparatuses. *Anopheles stephensi* mosquitoes have been mass-reared at the Radboudumc (Nijmegen, The Netherlands) since the 80s [3], with the insectary generating over 12,000 insects on a weekly basis for in- and external requests. Whilst the rearing protocol is highly standardized, a recently performed study evaluating the ingested blood meal volume after membrane feeding showed considerable variation between individual fully engorged mosquitoes [4], resulting in increased infection intensity heterogenicity due to the variable amounts of ingested parasites. We hypothesized that rearing procedures could be optimised to produce more homogeneous mosquito populations hence leading to more homogeneous blood meal volume and infection intensity.

Female *Anopheline* mosquitoes feed on blood from vertebrate hosts to acquire essential nutrients for reproduction. After oviposition, eggs typically hatch within 48 hours into motile larvae that cycle through four life-stages called instar (L1-L4), and then transform into nonfeeding pupae within which an adult mosquito develops and emerges from. Differences in egg hatching time and larval development directly influence mosquito body size, vectorial capacity and overall fitness [5–10]. The main factors determining these parameters are environmental conditions such as temperature and nutrition. Therefore, careful consideration is essential when selecting environmental conditions for laboratory-reared mosquitoes.

To explore possible optimizations of our standard mosquito rearing conditions, we compared the current mass-rearing procedure against an alternative approach, that aimed to optimize nutritional balance by standardizing the density of larvae per tray and tailored the diet to this larval density. We evaluated variation between protocols in consecutive mass-reared *Anopheline stephensi* mosquito generations by measuring body size and receptiveness to *P. falciparum* infection, using three indicators: wing-length, blood meal volume and infection intensity.

## Methods

### Collection and transferring mosquito eggs

*Anopheles stephensi* mosquitoes, Sind-Kasur strain originating from Pakistan [3] and in continuous culture at the Radboudumc since, were reared in a climate room at 30°C and ~80% relative humidity with a reverse light and dark cycle (12:12). Adult mosquitoes were maintained in cages (30 x 30 x 28 cm) made out of a plastic polymer bottom tray and a stainless steel frame that was covered with netting material. In order to obtain mosquito eggs, lithium heparinized blood (ref 367526, BD) from the national blood bank (Sanquin Nijmegen, The Netherlands) was fed using a maxi glass feeder connected to a circulating water bath set at 37°C [11]. Mosquitoes were allowed to feed for 30 min in the dark, and subsequently maintained on 5% glucose (D+ Glucose Monohydrate, ref 49159, Sigma) and 0.4% methylparaben (methyl 4-hydroxybenzoate, Sigma) at rearing conditions (day-0).

On day-3, a filter paper (Whatman) folded in a conical shape was placed in a disposable plastic cup (diameter 10 cm, high 5.5 cm) filled for 2/3 with tap water to keep it moist. The cup was kept in the cage overnight for the mosquitoes to lay eggs. Larval trays (HDPE plastic: 29 mm x 20.5 mm x 8 mm) were filled with tap water and left over night to be used the following day. On day-4 a small piece of filter paper (8 x 5 cm) with a hole cut-out in the middle was gently placed on the water surface of each tray, to prevent deposited eggs from dispersing and dry out. Subsequently, and based on experience, around ~400 eggs were transferred from the cup to the tray for the standard procedure by filter paper dipping (8 x 0.5 cm).

### Larval food

First instar larvae for the standard procedure were fed as described in **supplementary table 1**, using commercially available Liquifry (Liquifry No1, Interpet) and TetraMin baby (Tetra). To tip out the right amount of food for TetraMin baby, two holes, 2 mm (small) and 4 mm (large) of diameter, were made into the lid of the container

For the alternative food, first described by Damiens et al [12], bovine liver powder (Argentine Beef Liver Powder, NOW Foods) and tuna meal (LT Thunfischmehl, ref. Z30200, RSR-Baits) were used and first sieved to remove large particles. Subsequently, two parts of liver powder were added to two parts of tuna meal, and one part of vitamin mix (V1007-100G, Sigma Aldrich). Dry powder was thoroughly mixed and stored at 4°C for up to four weeks. Liquid food was prepared by dissolving 2% powder in deionized water followed by vigorously mixing by inversion. This suspension was stored at 4°C for up to three days and was mixed before use to avoid sedimentation. This custom-made larval food was transferred with a disposable pipette (ref 612, VWR) (**Supplementary table 2**).

### Collecting and maintaining mosquitoes

From day-9 onwards, trays were covered with nets to prevent emerged mosquitoes from escaping. Mosquitoes were collected daily with a custom made mobile electrical suction device (day-16 and −17) into separate cages (30 x 30 x 28 cm) and stored at 21°C and ~80% humidity. Each cage was provided with a 30 ml glass bottle filled with a solution of 5% glucose and 0.4% methylparaben (methyl 4-hydroxybenzoate, Sigma). A vertically rolled up Whatman filter paper was inserted to soak up the sugar solution and allow mosquitoes to feed on.

### Mosquito membrane feeding assay: blood meal volume, wing length and transmission

For feeding assays ~150 female mosquitoes (of similar age, between 1 and 3 days old) were transferred into midi-cages (20.5 x 15.5 x 19.5 cm), one hour before the membrane feeding assay was performed. For assessing malaria transmission, an infective blood meal with cultured *P. falciparum* gametocytes was prepared as described previously [13], and fed in duplicate (technical replicate). For blood volume analysis, 30 mosquitoes were fed heparin blood in cups (ref 367526, BD) from mini-feeders (volume of 300μL [11]), for 10 min in the dark. Upon visual inspection, unfed and partially fed mosquitoes were removed from the cage [11].

The blood meal volume from fully fed mosquitoes was analysed as described earlier [4]. For measuring wing length, mosquitoes were anesthetized with CO_2_ and right-wings were dissected in absence of fluid under a stereomicroscope. Wings were transferred with tweezers gently to glass slides and mounted by a coverslip. Length was measured from the alular notch to the axillary margin taking the second vein from above and excluding the wings fringes [14], by using an ocular micrometre (10:100, ref 474066-9901-000, Carl Zeiss). Malaria transmission intensity was assessed on day seven post infection, developed oocyst were counted by light microscopy (10x-40x magnification) after mercurochrome midgut staining (1%).

### Statistical analysis

Data was analysed with Graphpad Prism version 9. Difference in conditions were evaluated by t-test. Significance was indicated when value of *p* < 0.05 (\**p* < 0.05, \*\*\*\**p* < 0.001).

## Results

### Laboratory mosquito rearing procedure

Compared to our current rearing standard at the Radboudumc the alternative rearing procedure differed in water volume per tray, deposited eggs per tray (larvae density), and type of food. For practical reasons the rearing temperature was not evaluated in this study, and the standard climate room condition of 30°C and ~80% humidity was used. Before start of study a pilot for the alternative rearing procedure was conducted. With a water volume four times less per tray compared to the standard procedure evaporation was identified as a potential concern, as well as the exact amount of liquid food to provide per day/tray. Excess and insufficient amount of food negatively impacts larval developmental period with the latter ultimately resulting in mortality [9]. Water evaporation can worsen this, leading to an increased concentration of nutrients and waste products. During one cycle (14 days) we measured ~10% evaporation, therefore the volume per tray was increased to 700 ml to compensate for this loss.

Subsequently, the amount of food per day was assessed starting with 1 ml of liquid food for L1 larvae, and 2 ml for L2 larvae etc. In the standard protocol, a proximally 400 eggs were distributed per tray on day-4. For the alternative procedure this was 2500 eggs, but the following day exactly 200 larvae were counted and transferred with a disposable pipette (VWR, ref 612) to new trays. Mosquito development was assessed and visually compared to the trays where the standard rearing and feeding procedure was used (**Supplementary table 1**). It was observed that in the alternative procedure, the liquid food dissolved easily in water, excluding some larger particles that sank to the bottom of the tray. With a slightly adapted procedure to prevent over-feeding, we settled on a protocol that resulted in a reproducible pace of mosquito development (**Supplementary table 2**) and was used for the final comparison. The starting conditions for both procedures are summarized in **table 1**.

**Table 1:** *Anopheles stephensi* larvae starting conditions, for comparing standard breeding procedure against alternative. On day-3 larvae trays (HDPE plastic: 29 mm x 20.5 mm x 8 mm) were filled with tap water and at day-4 supplied with food before transfer of eggs. For the alternative procedure, 200 larvae were redistributed at day-5 to new trays, by manual counting using disposable pipettes. Following, larvae were fed as described in supplemental table 1 and 2.

|  |  | Standard | Alternative |
| --- | --- | --- | --- |
| Day-3 | Water volume per tray | 2,300 ml | 700 ml |
| Day-4 | Food per tray | Liquifry (1 drop) | 2 ml of liquid food |
| Day-4 | Eggs per tray | ~400 | ~2,500 |
| Day-5 | Larvae per tray | N/A | 200 exactly |

### Assessing the variation in body size and receptiveness to P. falciparum infection

To assess the possible optimization of the rearing conditions, the parental generation of female mosquitoes was blood fed and allowed to lay eggs at day-0, the start of study (experimental design in **Figure 1)**. Eggs were split four days later and reared according to the described procedures for three consecutive generations (**Supplementary table 1 and 2**). For generation 2 and 3, a subset of female mosquitoes of the same age (1 to 3 days old) was drawn from the stock cages and fed an uninfected or infectious blood meal for assessment of ingested blood volume or receptiveness to *P. falciparum* infection, respectively.

**Figure 1:**
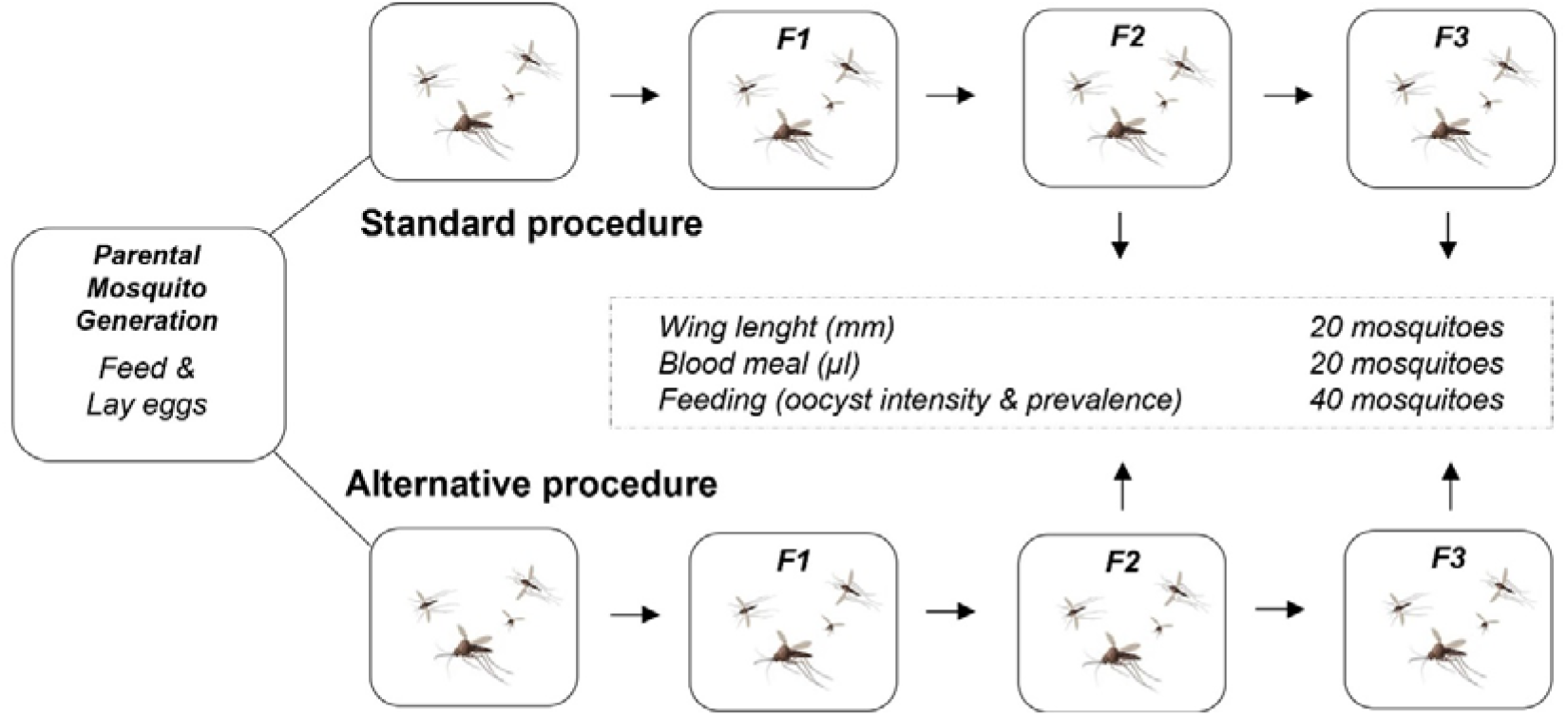
Experimental design for comparing mosquito breeding procedures. The standard rearing procedure was compared against the alternative procedure. Generation 2 and 3 mosquitoes were analysed for body size and receptiveness to *P. falciparum* infection.

The median mosquito wing length for generation 2 was 2.62 mm for the standard protocol (range 2.44-2.82, SD=0.08, n=20), and 2.47 mm for the alternative procedure (range 2.32-2.59, SD=0.07, n=20) (**Figure 2A**). For generation 3 this difference was confirmed: 2.68 mm for the standard protocol (range 2.53-2.82, SD=0.09, n=20), and 2.50 mm median mosquito wing length respectively (range 2.41-2.65, SD=0.07, n=20) (**Figure 2B)**. Mosquitoes reared with the standard procedure had in both generations significantly larger wings compared to the alternative procedure (*p* < 0.0001); indicating mosquitoes reared with this procedure were larger in size.

**Figure 2:**
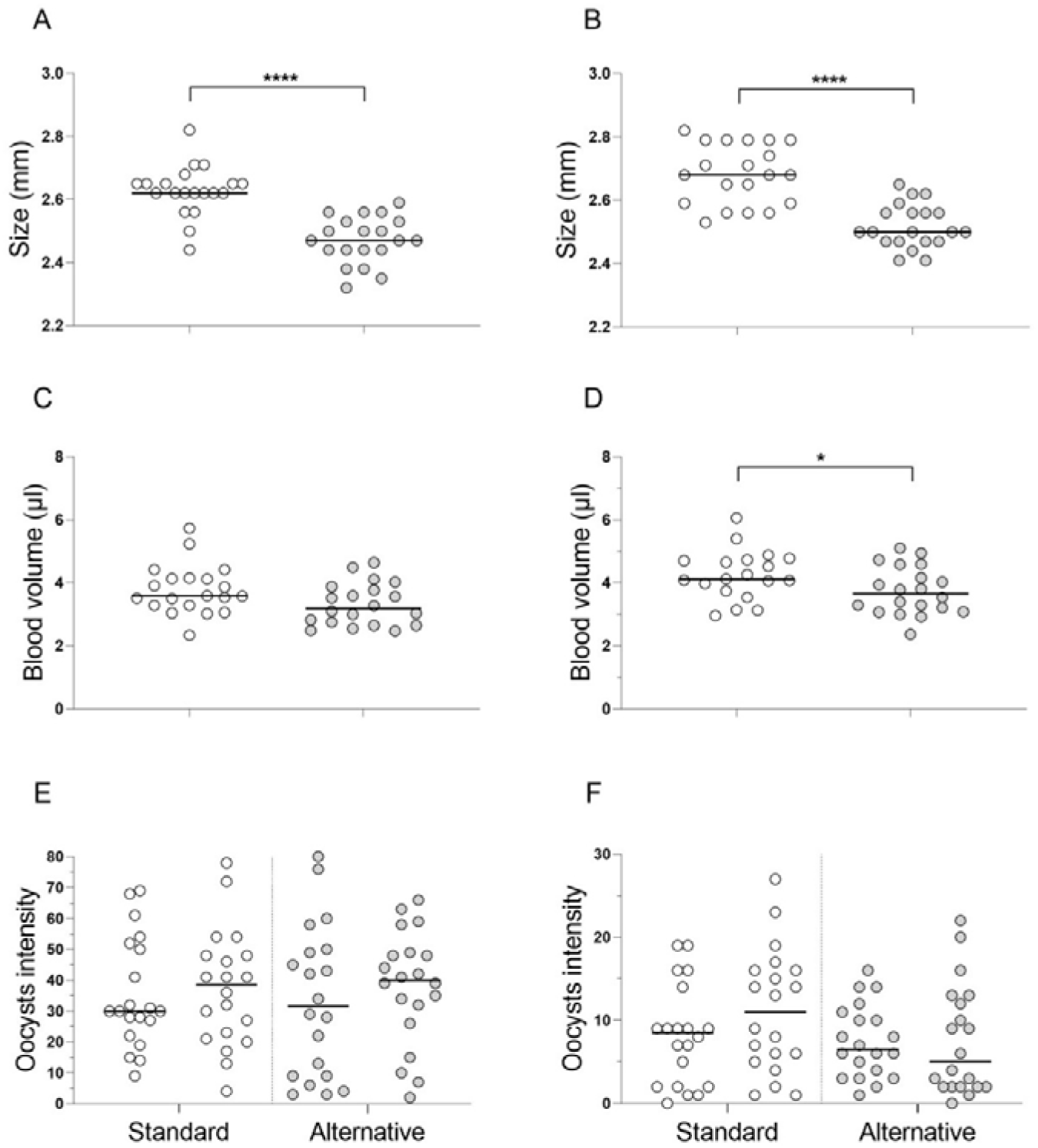
Comparative assessment of two *An. stephensi* laboratory rearing procedures. Mosquitoes were reared for three consecutive generations: F1-F3. Mosquitoes from generation 2 (panel **A**, **C** and **E**) and 3 (panel **B**, **D**, and **F**) were analysed to assess their body size and receptiveness to *P. falciparum*, by measuring the imbibed blood meal volume (μl) and wing-length (mm), or development of oocyst. Open circles indicate the standard, and closed circles the alternative procedure. Dots represent individual mosquitoes, and bars show the median. * *p* < 0.05, **** *p* < 0.0001.

For blood meal analysis, an alternative indicator for mosquito body size, fully engorged mosquitoes were processed directly after membrane feeding. The median imbibed volume for generation 2 was 3.59 μl (range 2.34-5.73, SD=0.78, n=20), and 3.19 μl in the alternative procedure (range 2.47-4.64, SD=0.67, n=20), *p*=0.0503 (**Figure 2C**). For generation 3 median 4.12 μl (range 2.96-6.06, SD=0.78, n=19) and 3.66 μl (range 2.37-5.1, SD=0.75, n=20), *p*=0.0452 (**Figure 2D)**. For both generations mosquitoes reared with the standard procedure imbibed a slightly larger volume of blood compared to the mosquitoes that were reared with the alternative procedure, but only in generation 3 these differences were statistically significant. Overall, both indicators for mosquito body size – wing length and imbibed blood volume – showed a similar trend, indicating mosquitoes reared with the standard procedure were larger in size.

Receptiveness to *P. falciparum* infection was subsequently assessed in duplicate (20 mosquitoes per feeder), seven days after feeding a homogenous infectious blood meal using NF54 *in vitro* cultured gametocytes. For generation 2, the median intensity was 30 and 38.5 oocysts for the standard procedure, and 31.5 and 40 oocysts for the alternative procedure, *p*=0.8403 (**Figure 2E**). For both procedures, prevalence of infection was 100%. For generation 3, the median intensity was slightly lower, presumably due to biological variation in infectivity of cultured gametocytes, with 8.5 and 11 oocysts in standard procedure compared to 6.5 and 5 oocysts for the alternative procedure, *p*=0.1214 (**Figure 2F**). Again, prevalence of infection was similar for both procedures (95%). For both generations, there was no statistically significant differences in median oocysts intensity between the standard and alternate procedure, and no difference in infection prevalence, suggesting that in these conditions the differences detected at the size level did not translate into infection yield differences.

## Discussion

This study aimed to explore possible optimizations of our standard *Anopheline stephensi* rearing procedure to generate more homogeneous mosquitoes. We assessed variation in body size and possible linkage to receptiveness to *P. falciparum* in consecutive mass-reared *Anopheline stephensi* mosquito generations, and compared our standard procedure to an alternative procedure with a lower density of larvae per tray and a different diet. For the alternative rearing procedure, the feeding regiment was optimized prior to start of study (**Supplementary table 2**), and there it was noted that this type of food easily dissolves. This may facilitate feeding but also made it difficult to visually assess how much food was needed per day. In contrast, the TetraMin baby food allowed for easy monitoring since it floated on the water surface and did not dissolve. However, once more familiar with the alternative feeding protocol for a given mosquito colony, food quantities per tray are likely to become more routine and the visual inspection of nutrient quantity thereby less important.

We further experienced that manual counting of larvae was time consuming, but there are automated methods available that in the future could improve the procedure [15, 16]. Assessment of two consecutive generations demonstrated a similar pace of mosquito development and no indications for changes in receptiveness to *P. falciparum* despite the larger body size of the standard protocol. Therefore, both protocols are suitable to apply in mosquito mass-rearing for assessing malaria transmission.

We hypothesized that alternative feeding procedure will lead to a larger and more homogenous mosquito population since this procedure involved a more controlled quantity of larval density and amount of food per tray [17]. Contrary to our expectations, mosquitoes reared with this procedure were slightly smaller in terms of wing-length and imbibed blood volume. Wing-length measurement showed a range of 0.38 mm between the smallest and largest mosquito in the standard procedure (generation 2 and 3 combined, 16% difference in length), but 0.33 mm in the alternative procedure (14%). The imbibed blood volume varied between mosquitoes but our average median of 3.85 μl for the standard protocol in this study (generation 2 and 3 combined, range 3.39, n=39), is very similar to what we previously reported for the same procedure (3.86 μl, range 4.1, n=94) [4]. For the alternative procedure this was lower, 3.42 μl (range 2.73, n=40). Variation between mosquitoes within one generation is natural and most likely caused by mosquito blood-feeding kinetics, such as the level of prediuresis (concentration of erythrocytes) [18], or to variation in body size. Because larvae in the alternative feeding procedure were collected/re-distributed one day later, the age of the larvae was likely to be more homogenous compared to the standard procedure. Interestingly, this did not result in a more synchronised pace of mosquito development (in our hands), with mosquitoes from both procedures emerging on the same day. Our findings do, however, suggest smaller variation in mosquito size with the alternative rearing method. This suggests that the alternative feeding protocol may not necessarily increase mosquito size but will reduce variation in size.

It should be noted that this work is by no means a comprehensive assessment of all factors related to optimal mosquito rearing. We therefore cautiously conclude that biological variation in size between mosquitoes seems inherent to mosquito biology development, and unavoidable in mosquito rearing. Interesting, the observed differences in wing size and engorged blood volume did not impact neither mosquito *P. falciparum* infection prevalence nor intensity. Further work is needed to determine the relevance of variation in bloodmeal volume in relation to heterogeneity in mosquito infection rates.

## Project funding statement

This work was funded by a European Research Council (ERC) Consolidator Grant to Teun Bousema [ERC-CoG 864180; QUANTUM], and the European Union’s Horizon 2020 Research and Innovation Programme under grant agreement no. 731060, Infravec2.

## Declarations

The authors declare that they do not have competing interests.

## Authors’ contributions

WG, GJ, GC, EAL, AY and TB, designed experiments; GJ, WG and KL performed laboratory rearing and infecting of mosquitoes; M.V-B and R.S. conducted gametocyte culture; GJ and WG performed mosquito dissection, and together with KL mosquito body size assessment; T.B. and WG analysed the data and drafted the first version of the manuscript. All authors read and approved the final manuscript.

## Acknowledgements

We would like to thank Astrid Pouwelsen, Jolanda Klaassen, Laura Pelser-Posthumus, Saskia Mulder and Jacqueline Kuhnen for their role in mosquito rearing and evaluating membrane feeding experiments. The authors are grateful to Jordache Ramjith for statistical analysis.

## Supplemental material

**Table 1:** Standard breeding and feeding procedure. Small tip, a hole of 2mm, large tip a hole of 4mm.

|  | Day | Week day | Activity | Feeding |
| --- | --- | --- | --- | --- |
| **Week 1** | 0 | Friday | Mosquito blood meal | - |
|  | 1 | Saturday | - | - |
|  | 2 | Sunday | - | - |
|  | 3 | Monday | Add cup with water to harvest eggs / Set up hatching trays with water | - |
|  | 4 | Tuesday | Transfer eggs to hatching trays | Liquify 1 drop |
|  | 5 | Wednesday |  | Small tip |
|  | 6 | Thursday |  | Small tip 2x |
| **Week 2** | 7 | Friday |  | Large tip |
|  | 8 | Saturday |  | - |
|  | 9 | Sunday | Apply netting | Large tip 2x |
|  | 10 | Monday |  | Large tip 3x |
|  | 11 | Tuesday |  | Large tip 4x |
|  | 12 | Wednesday |  | Large 4x |
|  | 13 | Thursday |  | Large 6x |
|  | 14 | Friday |  | Large 8x |
| **Week 3** | 15 | Saturday |  | - |
|  | 16 | Sunday | Collect mosquitoes | - |
|  | 17 | Monday | Collect mosquitoes | - |
|  | 18 | Tuesday | - | - |
|  | 19 | Wednesday | - | - |
|  | 20 | Thursday | - | - |
|  | 21 | Friday | Mosquito blood meal (day 0) | - |

**Table 2:** Alternative breeding procedure using custom-made liquid larvae food.

|  | Day | Week day | Activity | Feeding (volume) |
| --- | --- | --- | --- | --- |
| Week 1 | 0 | Friday | Mosquito blood meal | - |
|  | 1 | Saturday | - | - |
|  | 2 | Sunday | - | - |
|  | 3 | Monday | Add cup with water to harvest eggs / Set up hatching trays with water | - |
|  | 4 | Tuesday | Transfer eggs to hatching trays<br>Set up larvae trays with water | 2ml |
|  | 5 | Wednesday | Transfer larvae 200/tray | 2ml |
|  | 6 | Thursday |  | 1.5ml |
| Week 2 | 7 | Friday |  | 1.5ml |
|  | 8 | Saturday |  | - |
|  | 9 | Sunday | Apply netting | 2ml |
|  | 10 | Monday |  | 2ml |
|  | 11 | Tuesday |  | 2ml |
|  | 12 | Wednesday |  | 3ml |
|  | 13 | Thursday |  | 3ml |
|  | 14 | Friday |  | 3ml |
| Week 3 | 15 | Saturday |  | - |
|  | 16 | Sunday | Collect mosquitoes | - |
|  | 17 | Monday | Collect mosquitoes | - |
|  | 18 | Tuesday | - | - |
|  | 19 | Wednesday | - | - |
|  | 20 | Thursday | - | - |
|  | 21 | Friday | Mosquito blood meal (day 0) | - |

